# Human cerebral organoids capture the spatiotemporal complexity and disease dynamics of UBE3A

**DOI:** 10.1101/742213

**Authors:** Dilara Sen, Alexis Voulgaropoulos, Zuzana Drobna, Albert J. Keung

## Abstract

Human neurodevelopment and its associated diseases are complex and challenging to study. This has driven recent excitement for human cerebral organoids (hCOs) as research and screening tools. These models are steadily proving their utility; however, it remains unclear what limits they will face in recapitulating the complexities of neurodevelopment and disease. Here we show that their utility extends to key (epi)genetic and disease processes that are complex in space and time. Specifically, hCOs capture UBE3A’s dynamically imprinted expression and subcellular localization patterns. Furthermore, given UBE3A’s direct links to Angelman Syndrome and Autism Spectrum Disorder, we show that hCOs respond to candidate small molecule therapeutics. This work demonstrates that hCOs can provide important insights to focus the scope of mechanistic and therapeutic strategies including revealing difficult to access prenatal developmental time windows and cell types key to disease etiology.

## INTRODUCTION

Human cerebral organoids (hCO) are rapidly becoming important models for basic and translational research (Bagley et al., 2017; Lancaster et al., 2013; Di Lullo and Kriegstein, 2017; Pasca et al., 2015). Already, hCOs have been shown to accurately model the cell types and transcriptomics of early human neurodevelopment, which is particularly difficult to study due to the restricted availability of human fetal tissue (Camp et al., 2015; Li et al., 2017). They have also phenocopied disease processes including in ZIKA infection (Dang et al., 2016; Garcez et al., 2016; Qian et al., 2016), micro/lissencephaly (Bershteyn et al., 2017; Lancaster et al., 2013), Autism Spectrum Disorder (ASD) and related disorders (Birey et al., 2017; Mariani et al., 2015), and Alzheimer’s disease (Choi et al., 2016; Raja et al., 2016). Altered progenitor proliferation and death, excitatory-inhibitory imbalances, and aberrant gene expression were among the observable phenotypes using hCOs.

While this body of work is promising, the broader utility of hCOs in biomedical research will depend on their ability to recapitulate an increasing portfolio of neurodevelopmental details and phenotypes. In particular, neurodevelopment is inherently dynamic in space and time, with lessons from rodent models demonstrating that the timing, subcellular localization, and cell type specific expression of genes are indispensable for proper brain development (Mabb et al., 2011). It remains unclear to what extent hCOs accurately capture this level of spatiotemporal and epigenetic complexity. Further complicating matters, it remains unclear when in development the molecular underpinnings of neurodevelopmental disorders take root, and if these time periods even overlap those that hCOs can model (roughly the first half of gestation).

Of particular importance are the spatiotemporal dynamics of the over 100 imprinted genes that are allele-specifically expressed in the central nervous system (Bartolomei and Ferguson-Smith, 2011). Identifying where and when these alleles are expressed has deep implications for understanding the mechanistic etiologies of many neurodevelopmental disorders and for identifying potential time windows and cell types to target therapeutically. We decided to focus on *UBE3A*, an imprinted gene whose absence results in Angelman Syndrome (AS) and whose overabundance leads to a subset of ASD (Vatsa and Jana, 2018). We demonstrate that hCOs can model the complex spatiotemporal dynamics of UBE3A, reveal dynamic phenotypes specific to humans, and respond to candidate small molecule therapeutics. The results of this work provide evidence to motivate the broader use of hCOs to investigate imprinted loci, other complex epigenetic phenomena, and their related neurodevelopmental disorders.

*UBE3A* is an ideal candidate to test the utility of hCOs because of the existence of strong pioneering work in mouse models (Lopez et al., 2019). These provide important biological context and molecular benchmarks for hCOs. There are also significant anatomical differences between mouse and human brains including in UBE3A isoforms (LaSalle et al., 2015) and cis-regulatory elements (Chamberlain et al., 2010; Hsiao et al., 2019) that could result in substantially different *UBE3A* biology. Thus, hCOs also present an opportunity to identify key species-specific differences, with both similarities and differences potentially providing key insights into disease mechanisms and treatment strategies. Thus, the overall strategy in this work is to harness salient features of mouse *UBE3A* as benchmarks for hCOs, while also identifying human-specific *UBE3A* properties. The key questions we asked were: 1) are there specific time periods during which human UBE3A is imprinted or its subcellular localization changes that indicate potential therapeutic time windows, and do hCOs even capture these time periods; 2) in which human cell types does UBE3A exhibit nuclear localization, which was recently functionally connected to disease phenotypes (Avagliano Trezza et al., 2019); and 3) can hCOs be used to test or screen for potential therapeutic compounds?

## RESULTS

### hCOs reveal an early transition of cytoplasmic to nuclear UBE3A

One of UBE3A’s salient molecular features is its shift from the cytoplasm to nucleus upon neuronal maturation (Burette et al., 2017; Dindot et al., 2008; Gustin et al., 2010), likely driven by shifts in isoform expression (LaSalle et al., 2015; Miao et al., 2013; Sadhwani et al., 2018). Importantly, it was recently shown that mice lacking nuclear UBE3A show electrophysiological and behavioral deficits similar to other AS model mice that lack maternal *UBE3A* (Avagliano Trezza et al., 2019). Furthermore, UBE3A has two putative functions, regulating gene expression and protein degradation (LaSalle et al., 2015), that could be influenced by subcellular localization. Thus, mapping UBE3A’s subcellular localization over time and across cells types is important for understanding disease etiology and treatment.

Given the importance of UBE3A’s subcellular localization, we asked if ‘whole brain’ hCOs (Lancaster et al., 2013) would recapitulate this nuclear shift, and furthermore, if hCOs could be used to efficiently map when and in which cell types *UBE3A* is expressed. As a benchmark to orient our experiments, prior work showed that just after birth (i.e. P0-6 or ~3 weeks post conception), mouse UBE3A transitioned from diffuse to strongly nuclear in many neurons of the brain (Judson et al., 2014). hCOs have previously been shown to model cell types up through the first human trimester by comparisons to human fetal tissue (Camp et al., 2015; Luo et al., 2016; Quadrato et al., 2017), but have yet to be grown to the equivalent of postnatal maturity. Furthermore, there is a lack of data on the prenatal biology of UBE3A. Thus, we were unsure if the nuclear localization of UBE3A seen postnatally in mice would be observable in hCOs. However, the timing of mouse and human neurodevelopment is difficult to directly correlate with some aspects reliant on absolute timescales (Mason and Price, 2016). Therefore, we grew, fixed, and sectioned hCOs over a broad time range (1-12 weeks) in culture and stained for nuclei, UBE3A, and a diverse panel of cell-type specific markers (Figure 1, S1, 2, and S2). Interestingly, we found that UBE3A localized strongly in the nuclei of a substantial number of neurons after only 3 weeks in culture, matching or even preceding mice on an absolute timescale (Figure 1A and S1A). The percentage of cells with nuclear UBE3A increased over time, correlating with the appearance of PAX6+ neural progenitors and TUJ1+/SOX2− neurons (Figure 1B, 1C, and S1). In fact, the lineage progression of these three cell types seemed to track the shift in UBE3A localization: SOX2+ stem cells showed almost no nuclear UBE3A, roughly half of PAX6+ progenitors exhibited strong nuclear UBE3A, and TUJ1+ neuronal areas showed almost complete nuclear staining (Figure 1D). Thus, human UBE3A recapitulates the same general nuclear localization in neurons as in mice and at a similar absolute timescale, but at a much earlier stage relative to birth. Furthermore, hCOs appear to capture the important time periods to study and observe UBE3A dynamics.

**Figure 1.**
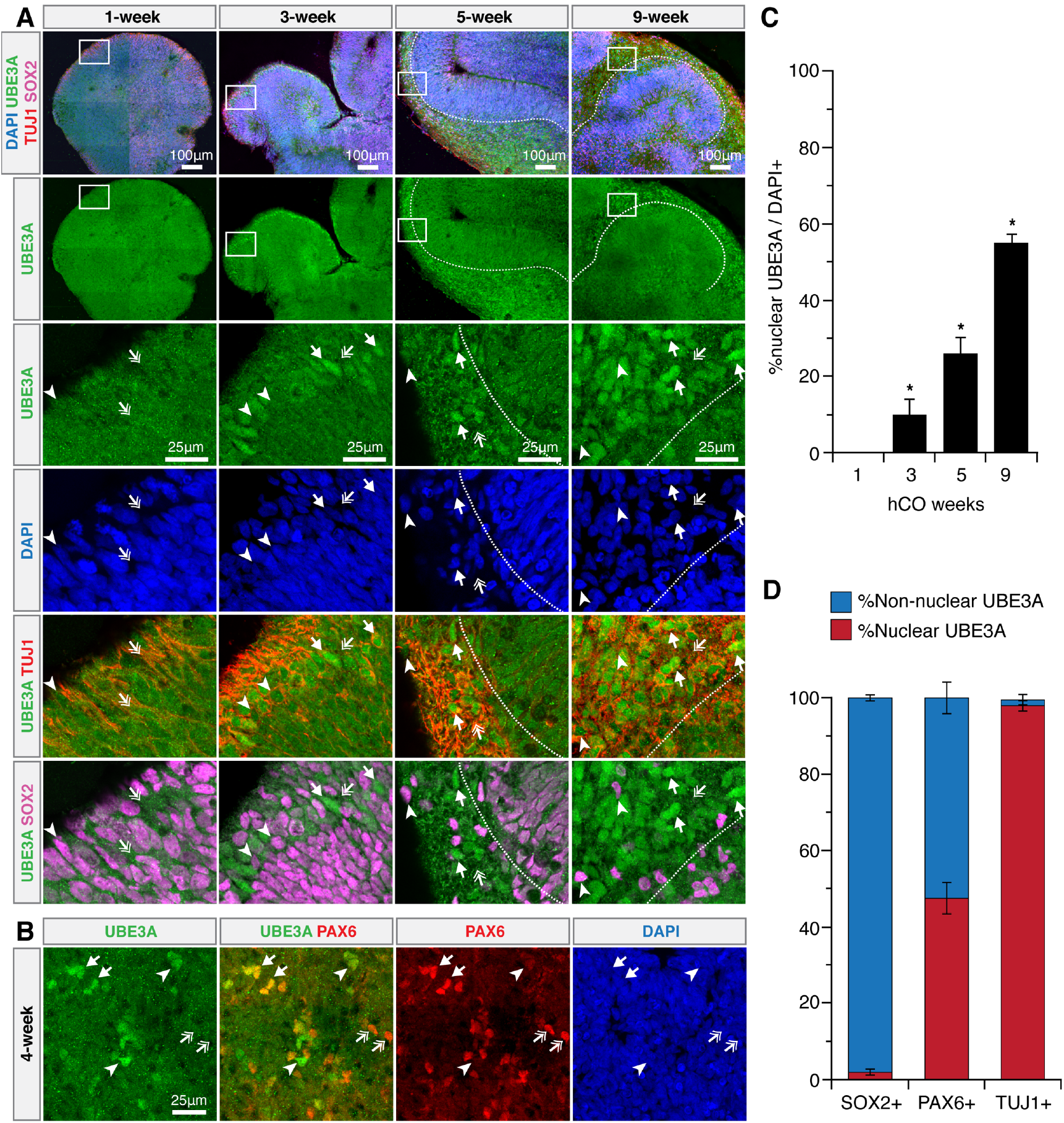
hCOs reveal an early transition of cytoplasmic to nuclear UBE3A. (A) Immunostaining time course of neurotypical hCO neurodevelopment. (A) Boxes bound high magnification images. Nuclear UBE3A in neurons (arrows). Diffuse UBE3A in SOX2+ cells (arrow heads). Decreasing cytoplasmic UBE3A over time (double arrows). Dotted white lines delineate boundaries between TUJ1+ and SOX2+ cells. (B) Nuclear (arrows) and diffuse (double arrows) UBE3A in PAX6+ cells. Nuclear UBE3A in PAX6-/weak cells (arrow heads). (C) % nuclear UBE3A increases during hCO development. Immunostaining quantification. *P<0.05 between all groups, one-way ANOVA with Tukey-Kramer post hoc analysis, n=3-6 hCOs per time point. Error bars = 95% confidence intervals. (D) UBE3A localization by cell type. Immunostaining quantification. Error bars = 95% confidence intervals.

**Figure 2.**
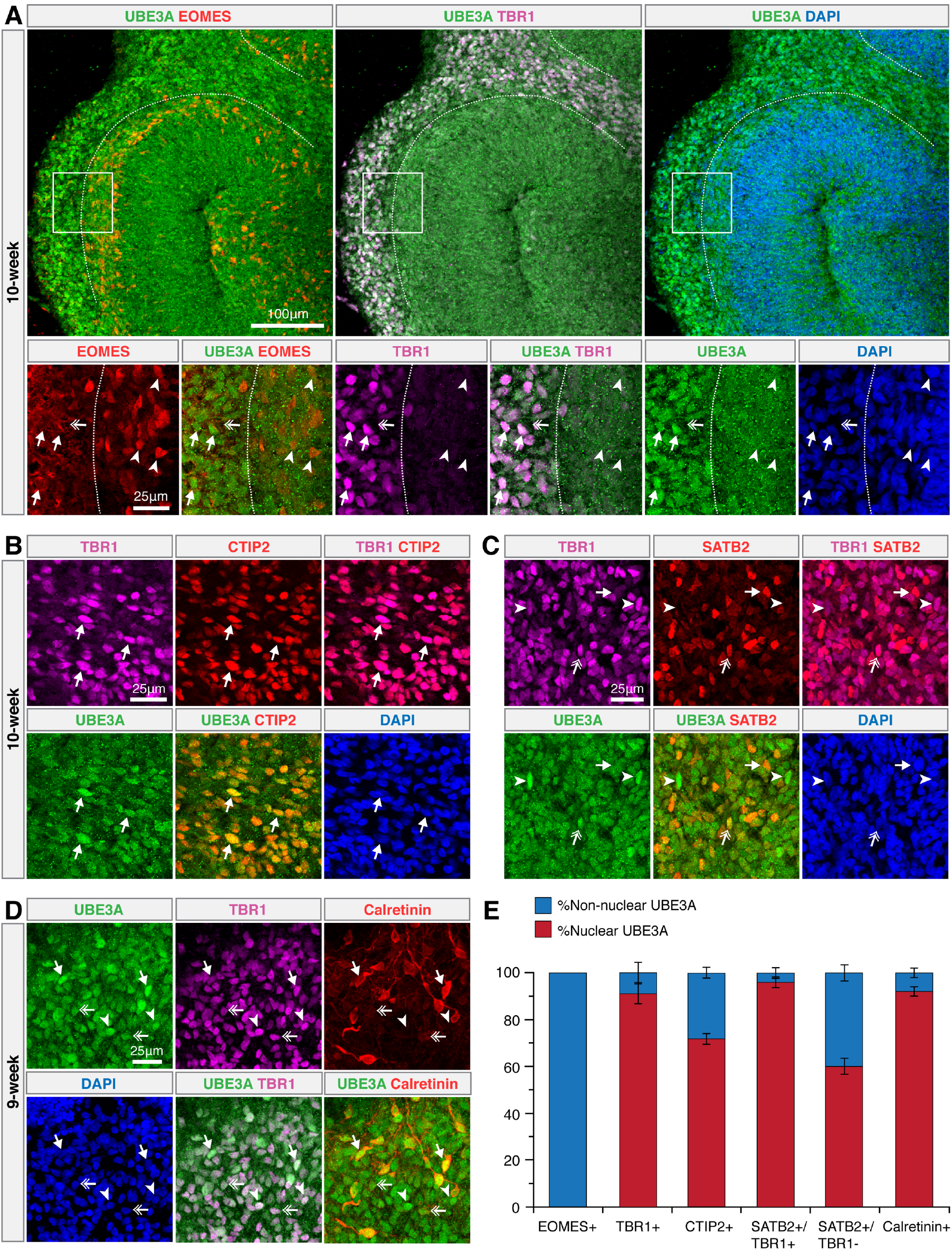
Cortex-like development in hCOs provides temporal benchmarks for UBE3A dynamics. (A) Dotted white lines delineate boundaries between TBR1+ and EOMES+ regions. Boxes bound high magnification images. Nuclear UBE3A in TBR1+ cells (arrows). Diffuse UBE3A in EOMES+ cells (arrow heads). Weak UBE3A signal in cytoplasm of TBR1+ cells (double arrows). (B) Nuclear UBE3A colocalizes with CTIP2+/TBR1+ cells. (C) Strong nuclear UBE3A in TBR1+/SATB2− (arrow heads) and TBR1+/SATB2+ (double arrows) cells. Diffuse UBE3A in TBR1−/SATB2+ cells (arrows). (D) Nuclear UBE3A in TBR1+/Calretinin+ (arrows), TBR1+/CalretininWeak (double arrows) and TBR+/Calretinin- (arrow heads) neurons. (E) UBE3A localization by cortical cell type. Immunostaining quantification. Error bars = 95% confidence intervals. n=3-6 hCOs.

### Cortex-like development in hCOs provides temporal benchmarks for UBE3A dynamics

Given the difficulty of correlating absolute and gestation-relative developmental timescales between species and artificial systems such as hCOs, we next asked if there is a structural boundary that reflects developmental progression that could be used to assess UBE3A dynamics across species. The fetal cortex is comprised of layers of cells representing different stages of differentiation, including a sequential transition from neural precursors (SOX2), to radial glia and intermediate progenitors (PAX6, EOMES) to post-mitotic neurons (TBR1, CTIP2, SATB2) (Englund, 2005). Interestingly, in 10 week old hCOs, we observed a striking boundary between cortex-like layers of TBR1+ and EOMES+ cells in hCOs, with strong nuclear UBE3A only in TBR1+ cells (Figure 2A). As TBR1+ pre-plate regions are precursors to the cortical-plate we stained for CTIP2 (early-born deep layer neurons) and SATB2 (late-born superficial layer neurons) in 10-week old hCOs and found that both markers were expressed in cells with strong nuclear UBE3A (Figure 2B, S2A, S2B and S2C). Furthermore, while relatively rare, mature SATB2+/TBR1− cells exhibited a relative loss in nuclear UBE3A compared to their more common and immature SATB2+/TBR1+ counterparts (Figure 2E). Interestingly, the strong coexpression of TBR1 with SATB2 and CTIP2 cells correlates with periods before human post conception week 20 (PCW20), with separation of these markers occurring closer to GW30 in human fetal tissue (Saito et al., 2011), suggesting that these 10 week hCOs may be roughly capturing neurodevelopmental periods before GW20.

As another method to infer the “analogous gestational age” of when UBE3A is localizing to the nucleus in hCOs, we analyzed the co-expression of calretinin with TBR1. It was previously reported that calretinin expression in the early fetal brain is not restricted to GABAergic interneurons but is also present in the first excitatory projection neurons of the cortex (labeled by TBR1). During human PCW7-7.5, TBR1+/calretinin+ cells co-exist with TBR1−/calretinin+ neurons in the advanced pre-plate and the pioneer cortical-plate. Shortly after this stage, the cortical plate is established (PCW8) where TBR1+/calretinin- and TBR1+/weak calretinin neurons are observed (Gonzalez-Gomez and Meyer, 2014). This led us to the idea that the co-expression of calretinin with TBR1 and subsequent weakening and disappearance of calretinin may serve as a pseudo-temporal benchmark for hCOs. Indeed, in 9 week old hCOs we were able to capture this transition point where both TBR1+/calretinin+ and TBR1+/weak calretinin neurons were observed (Figure 2D and S2D). In both of these cell types UBE3A was localized primarily to the nucleus and the expression of UBE3A seemed to be higher in calretinin+ neurons (Figure 2D and S2D). Collectively these results suggest the nuclear localization of UBE3A in hCOs occurs in what could be the equivalent of the 1^st^ trimester of human gestation. In addition, this strategy could be more broadly applied in the future to use key features of cortical development to benchmark other molecular events in hCOs to a rough gestational timeline.

### hCOs reveal distinct patterns of UBE3A expression in progenitor-related compartments

A significant advantage of hCOs is their ability to create many cell types from the whole brain. Given the functional importance of UBE3A localization (Avagliano Trezza et al., 2019), and the ability of hCOs to capture this localization (Figures 1, S1, 2, S2), we asked if there were unique expression patterns in cell types other than neurons that have yet to be observed. UBE3A was previously found not to imprint in mouse stem cell and progenitor compartments (Rougeulle et al., 1997; Yamasaki et al., 2003) and there is growing evidence that altered progenitor properties may contribute to AS (and ASD) phenotypes including microcephaly and delayed myelination (Judson et al., 2017; Singhmar and Kumar, 2011). Thus, we focused on the cortical progenitor niche and the choroid plexus, which generates cerebral spinal fluid and its associated factors that influence niche properties (Huang et al., 2010; Johansson et al., 2013).

We observed two unique staining patterns not previously reported in mouse or human systems. First, we observed a relative absence of UBE3A in the nuclei of phospho-vimentin+/SOX2+ progenitors, and a high intensity perinuclear UBE3A signal instead (Figure S2E). Progenitors from the apical-like surface and ventricular/sub-ventricular zone-like regions both exhibited these expression patterns. Second, we identified a short time period between 3-5 week old hCOs where UBE3A was strongly nuclear in the TTR+ (transthyretin) epithelial cells of choroid plexus-like regions (Figure S2F). This pattern was lost in older hCOs (7 weeks) with UBE3A staining more diffuse across subcellular compartments. While it is unclear if these specific spatiotemporal patterns indicate biologically relevant functions of UBE3A, they suggest an intriguing possibility that neurogenesis impacts AS/ASD disease etiology.

### hCOs recapitulate the complex epigenetic expression of *UBE3A* and *UBE3A-ATS*

While UBE3A’s subcellular localization is important for its function, its dosage is also a clear primary driver of AS and ASD disease phenotypes. As an imprinted gene, *UBE3A’s* paternal allele is silenced just after birth in mice (between P0-P6) in most neurons of the brain (Judson et al., 2014). Because the paternal allele is silenced, mutations or deletions of the maternal allele result in the absence of UBE3A protein in neurons, resulting in Angelman Syndrome (Kishino et al., 1997). Conversely, maternal *UBE3A* duplications and gain of function mutations are linked to ASD phenotypes (Yi et al., 2017). The silencing of paternal *UBE3A* is also associated with a long, non-coding antisense RNA (*UBE3A-ATS).* As cells differentiate into neurons, *UBE3A-ATS* expression increases to a high enough level to silence paternal *UBE3A*, potentially through a hypothesized collision mechanism (Hsiao et al., 2019; Stanurova et al., 2016) between these two opposed and overlapping transcripts.

Given the importance of *UBE3A-ATS* and *UBE3A*, we asked whether their dynamic epigenetic expression could be captured in hCOs. Through quantitative RT-PCR, we observed a monotonic increase in *UBE3A-ATS* transcripts starting at 3 weeks in culture (Figure 3A). However, we could not observe paternal *UBE3A* silencing in neurotypical hCOs (Figure 3B). This is expected as previous reports in mice showed that maternal *UBE3A* dosage compensates for paternal imprinting (Hillman et al., 2017). Therefore, to observe paternal *UBE3A* silencing we obtained a human induced pluripotent stem cell line previously generated from an Angelman syndrome patient (Chamberlain et al., 2010) and created hCOs from them (AS hCO). These cells lack maternal *UBE3A* so any transcripts detected would be derived from the paternal copy. When we measured *UBE3A-ATS* transcript levels, we observed a monotonic increase, similar to neurotypical hCOs (Figure 3A). As expected, *UBE3A* transcripts decreased only in AS hCOs, after 6 weeks (Figure 3B). In addition, the ~3 week delay between when *UBE3A-ATS* began to increase and *UBE3A* decreased is similar to previous observations (Hsiao et al., 2019). We also observed an interesting transient increase in *UBE3A* in both neurotypical and AS hCOs, suggesting that a transient elevation in *UBE3A* expression may occur in neurodevelopment prior to imprinting.

**Figure 3.**
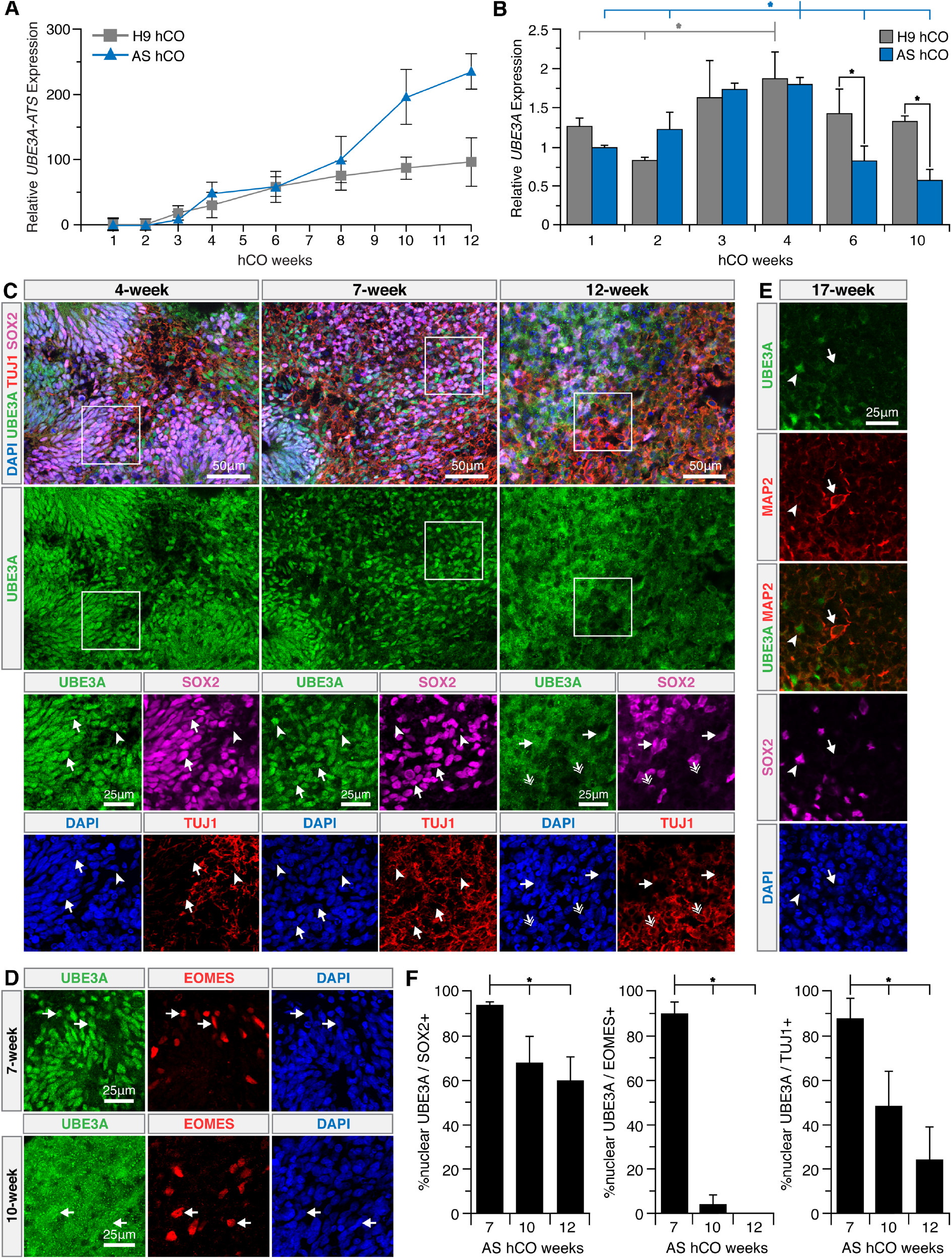
hCOs recapitulate the complex epigenetic expression of *UBE3A* and *UBE3A-ATS*. (A and B) QRT-PCR measurements of mRNA levels of *UBE3A-ATS* (A) and *UBE3A* (B) in neurotypical and AS hCOs, normalized to HPRT, ratioed to 1-week-old AS hCOs. Error bars are 95% confidence intervals. (A) p<0.05 t-test against null slope hypothesis, n=3 biological replicates. (B) *p<0.05, full tick marks compared to half tick marks by one-way ANOVA with Tukey-Kramer post hoc, n=3 biological replicates. (C-E) UBE3A expression and localization in time course of AS hCOs. (C) Boxes bound high magnification images. Nuclear UBE3A in SOX2+ progenitors of 4-12 week old AS hCOs (arrows). Nuclear UBE3A in 4-7 week old TUJ1+/SOX2− neurons (arrow heads) is lost in 12 week old AS hCOs (double arrows). (D) Nuclear UBE3A in 7 week old EOMES+ cells (arrows) is lost at 10-weeks in AS hCOs. (E) UBE3A is absent in 17 week old MAP2+/SOX2− neurons (arrows). SOX2+ progenitors still express some paternal UBE3A (arrow heads). (F) % nuclear UBE3A in 7-12 week old AS hCOs. Immunostaining quantification. *P<0.05, full tick marks compared to half tick marks by one-way ANOVA with Tukey-Kramer post hoc analysis, n=3-6 biological replicates. Error bars are 95% confidence intervals.

Taking further advantage of the fact that AS hCOs have only paternal *UBE3A* we next asked where paternal UBE3A was being localized over time. Interestingly, unlike in neurotypical hCOs, UBE3A localized to the nucleus of SOX2+ and EOMES+ progenitors during early hCO development (4-7 weeks) (Figure 3C, 3D, S3A). We suspect that the apparent nuclear localization of UBE3A in progenitors could be due to the absence of cytoplasmic isoforms, although the strength of expression was similar across neurons and neural progenitors. In older AS hCOs (10-12 weeks), UBE3A expression became substantially more diffuse in EOMES+ cells (Figure 3D, 3F and S3B), while becoming only slightly less nuclear in SOX2+ cells (Figure 3F). In neurons of 1-7 week old hCOs, UBE3A was nuclear, but upon extended culture (10-17 weeks) UBE3A intensity weakened, becoming absent in TUJ1+, MAP2+, TBR1+ and SATB2+ neurons (Figure 3C, 3E, 3F, S3A, and S3C-E). Interestingly, immature SOX2+/TUJ1+ neurons did exhibit nuclear UBE3A in 10-12-week old AS hCOs, consistent with previous reports of paternal UBE3A expression in immature neurons (Grier et al., 2015; Jones et al., 2016; Judson et al., 2014; Sato and Stryker, 2010) (Figure S3F). Collectively, these results suggest hCOs exhibit successful imprinting and that this process may be occurring very early in human fetal development.

### Paternal *UBE3A* can be pharmacologically re-activated in AS hCOs

Since AS hCOs successfully imprint paternal *UBE3A* and model early human neurodevelopment, they represent a useful new experimental system to test and screen potential AS therapeutics. A leading therapeutic strategy actively being developed for Angelman Syndrome has been to reactivate the imprinted paternal *UBE3A* to compensate for the absent maternal copy (Bailus et al., 2016; Chamberlain and Brannan, 2001; Huang et al., 2012; Lee et al., 2018; Meng et al., 2013, 2014). We therefore asked if paternal *UBE3A* could be pharmacologically reactivated in AS hCOs. Prior work found topoisomerase inhibitors (topotecan and indotecan) could reactivate paternal *UBE3A* in mice (Huang et al., 2012; Lee et al., 2018) and human cell cultures (Fink et al., 2017; King et al., 2013). In particular, these compounds were shown to reactivate paternal *UBE3A* by decreasing paternal *UBE3A-ATS* levels. Therefore, we first used 11-week-old hCOs as they showed significant imprinting of *UBE3A*, and added 1 μM topotecan or indotecan and measured *UBE3A-ATS* and *UBE3A* transcripts 3 days after treatment. Both topotecan and indotecan were able to knock down *UBE3A-ATS* 7 and 4 fold while increasing *UBE3A* 1.8 and 1.75 fold, respectively (Figure 4A). These roughly match changes observed in mouse and human neuronal culture systems(Fink et al., 2017; Huang et al., 2012). Furthermore, we observed an increase in nuclear UBE3A in neurons through immunostaining (Figure 4B, 4C, and S4B).

**Figure 4.**
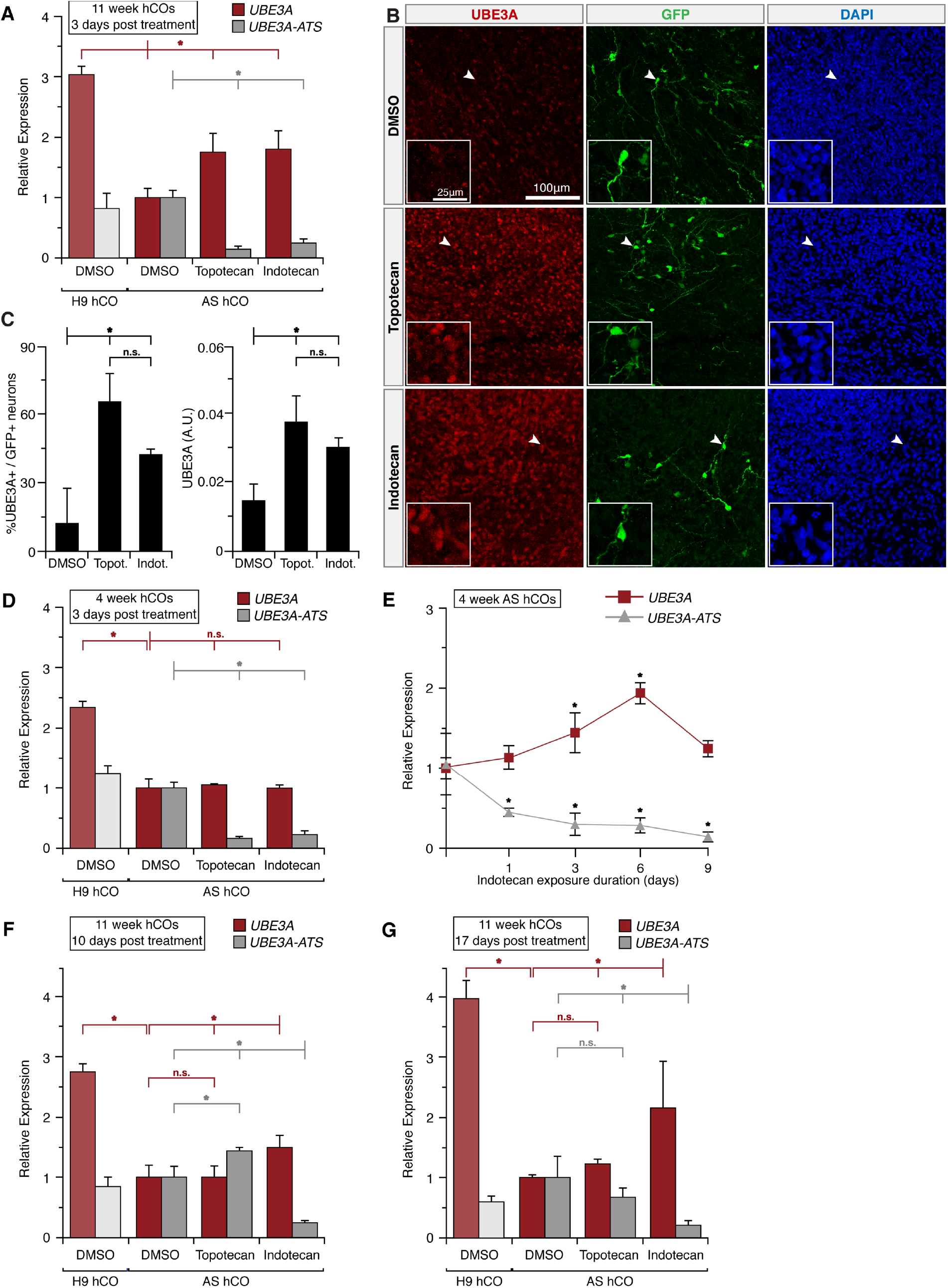
Paternal *UBE3A* can be pharmacologically re-activated in AS hCOs. (A, D, F, G) QRT-PCR measurements of mRNA levels of *UBE3A* (red) and *UBE3A-ATS* (grey) after vehicle (DMSO), 1μM Topotecan, or 1μM Indotecan treatment. Signals normalized to TBP and ratioed to vehicle treated AS hCOs. *P<0.05, n.s. not significant, full tick marks compared to half tick marks by one-way ANOVA with Tukey-Kramer post hoc, n=3 biological replicates, error bars = 95% confidence intervals. A.U. arbitrary fluorescence units. (A) mRNA 3 days after a single drug-treatment in 11 week hCOs. (B) 11 week old AS hCOs with CamKIIa-GFP neurons. Insets zoom in on arrow heads. (C) Immunostaining quantification of (B). (D) mRNA 3 days after a single drug-treatment in 4 week old hCOs. (E) mRNA after 1-9 days of continuous 1 μM indotecan treatment in 11 week old AS hCOs, ratioed to untreated day 0 AS hCOs. No significant change in vehicle treated samples (Figure S4C). *P<0.05, compared to day 0 by one-way ANOVA with Tukey-Kramer post hoc, n=3 biological replicates, error bars = 95% confidence intervals. (F) mRNA 10 days after a single drug-treatment in 11 week hCOs. (G) mRNA 17 days after a single drug-treatment in 11 week hCOs.

One the major open questions in treating neurodevelopmental disorders is the delivery kinetics and pharmacodynamics. At what time point, how frequently, and for how long should potential therapeutics be delivered, and how persistent are therapeutic effects? We therefore asked if topotecan/indotecan treatments would have differential effects if delivered to different age AS hCOs (4, 11, and 15 weeks). Both compounds knocked down *UBE3A-ATS* and increased *UBE3A* at 11 and 15 weeks (Figure 4A and S4A). However, at 4 weeks, we observed only a reduction in *UBE3A-ATS* and did not detect an increase in *UBE3A* expression (Figure 4D). This could be attributed to a delay between transcription of *UBE3A-ATS* and silencing of *UBE3A* (Hsiao et al., 2019; Stanurova et al., 2016) (Figures 3A and 3B) so that *UBE3A* expression levels are still high at that stage of hCO development.

Next, we asked if the rescue of *UBE3A* could be enhanced by persistent and longer-term therapeutic delivery. We exposed AS hCOs to fresh media prepared with 1 μM indotecan every day for 9 days and analyzed samples at days 1, 3, 6, and 9. Our results showed that *UBE3A* levels increased up to day 6. However, the samples analyzed on day 9 showed a decrease in *UBE3A* expression. *UBE3A-ATS* levels remained low throughout this analysis with no significant differences between time points (Figure 4E and S4C).

The decrease in *UBE3A* after 9 days of repeated treatments may have been due to the toxic effects of topoisomerase inhibitors (Kummar et al., 2016; Lee et al., 2018). We therefore asked if a single pharmacological treatment could have lasting effects. We delivered topotecan or indotecan to 11 week old AS hCOs and measured transcript levels 10 and 17 days later with media changes containing no drugs after the single treatment. While topotecan was unable to maintain elevated levels of *UBE3A* after both 10 and 17 days, indotecan’s rescue of *UBE3A* persisted throughout the experiment (Figure 4F and 4G). The increased ‘memory’ of indotecan could be due to its increased chemical stability (Kummar et al., 2016; Lee et al., 2018) or an as yet unknown epigenetic mechanism. We also note that topotecan treated samples appeared to have a compensatory rebound 10 days after treatment in which *UBE3A-ATS* increased beyond the original levels (Figure 4F). In addition, both topotecan and indotecan were unable to fully rescue *UBE3A* to neurotypical levels, although studies suggest even partial rescue of *UBE3A* would have beneficial effects (Brockmann et al., 2002; Le Fevre et al., 2017; Meng et al., 2012). Collectively, these observations may inform the treatment window and regimen for potential future therapeutics.

## DISCUSSION

The excitement surrounding hCOs derives from their potential to fill important gaps that are inaccessible by other experimental systems. This is illustrated in this work, where important molecular events involving *UBE3A* were found to be dynamically occurring early in hCO development, supporting the hypothesis that the etiology of many Angelman Syndrome phenotypes may arise prenatally (Silva-Santos et al., 2015; Sonzogni et al., 2019). These results emphasize a primary utility of hCOs: they provide ample physiologically relevant human materials to access prenatal time periods that are difficult to obtain even in animal models and that face ethical and legal restrictions in humans. hCOs may also serve as experimental models to ‘translate’ analogous phenotypes and their timescales observed in animal models to human biology; for example, similar molecular events such as a change in protein localization or imprinting could be compared across the very different gestational timescales of mice and humans. This work presents a broadly applicable archetype for how hCOs can be harnessed to access these gaps in our understanding of human neurodevelopment and associated disorders, and how they relate to and could potentially leverage useful and longstanding animal models.

Another major advantage of hCOs is their ability to generate a diverse range of human cell types at very early points in neurodevelopment. For example, we showed that the subcellular localization of UBE3A could be tracked in time but also across diverse cell types present during neurodevelopment. We further demonstrated the multifold importance of this capability: it identifies prenatal stages that may be important for disease etiology; it hints at effective time windows for administering therapeutics; and it suggests roles for specific cell types in disease mechanisms, including cell types not present in rodent models (i.e. outer radial glia). For example, the distinct localization patterns of UBE3A in stem cells, progenitors, and choroid plexus-like cells that we observed (Figures 1B, S1B, 2A and S2E and S2F), including differential expression patterns between neurotypical and AS hCOs (Figures 3C-F), were previously not known; but they strongly suggest future work should investigate the mechanistic roles of these cell types in AS and ASD.

As with subcellular localization, imprinting is also a dynamic feature crucial in neurodevelopment. Imprinting affects gene dosage, potentially mediates environmental influences (Kappil et al., 2015; Litzky et al., 2018), and its misregulation is implicated in a range of neurodevelopmental disorders (LaSalle et al., 2015). Given that the brain is host to over 100 tissue-specific imprinted genes, the utility of hCOs could be quite broad. It should be noted that especially with regards to imprinted genes, human cell lines should be chosen carefully. The imprinted status of the Prader-Willi Syndrome imprinting center, relevant to *UBE3A* imprinting, had previously been confirmed for both the H9 and AG1-0 pluripotent cell lines used in this study (Chamberlain et al., 2010; Stanurova et al., 2016). In addition, *UBE3A* expression was previously shown to imprint during neuronal differentiation of these cells in 2D cultures. However, a recent large scale study of over 270 hPSC lines found that while most cell lines maintained proper imprinting at most loci, the majority had at least one locus that lost proper imprinting (Bar et al., 2017). Thus, specific hIPSCs may not be suitable for studies involving certain loci.

Perhaps the most important aspect of *UBE3A* biology revealed in this work is the fact that both the salient changes in subcellular localization and in *UBE3A-ATS/UBE3A* expression occurred in what is the hCO equivalent of the first human trimester. The potential implications of these early dynamics are profound. While hCOs cannot provide behavioral phenotypes, recent work from Elgersma and colleagues have shown, through conditional control of *UBE3A* knockout or reinstatement, that at least a subset of behaviors in mice are impacted by perinatal *UBE3A* levels and cannot be rescued later in neurodevelopment (Rotaru et al., 2018; Silva-Santos et al., 2015; Sonzogni et al., 2019). Collectively, both human and mouse studies now support a scenario where early, even prenatal, treatment in humans may be necessary to have maximal therapeutic effects, although significant benefits may still be achieved through interventions later in life. In addition, hCOs may help inform specific delivery schedules. For example, we found: early treatment prior to *UBE3A* imprinting would likely be ineffective (Figure 4D); indotecan may have more persistent effects than topotecan (Figure 4F and 4G); and transient administration of some therapeutics may lead to an undesired rebound effect that should be carefully considered or monitored (Figure 4F).

hCOs have advanced rapidly with studies demonstrating their similarity to fetal tissue (Camp et al., 2015; Quadrato et al., 2017), protocols that create hCOs with remarkably consistency (Velasco et al., 2019), and even modularly combined hCOs (Bagley et al., 2017; Giandomenico et al., 2019; Xiang et al., 2017). Here we have shown that the utility of hCOs can be expanded in the contexts of tracking complex spatiotemporal and epigenetic molecular events, identifying new molecular phenotypes in diverse cell types, and modeling therapeutically relevant responses to distinct treatment regimens.

## ACKNOWLEDGEMENTS

This work was supported by a Simons Foundation SFARI Explorers Grant (495112), the NCSU Faculty Research and Professional Development Program, an NCSU Research Innovation Seed Fund Grant, the NSF Emerging Frontiers in Research and Innovation program (NSF-1830910), an NIH Avenir Award (DP1-DA044359), and a fellowship from the American Association of University Women (to DS).

## AUTHOR CONTRIBUTIONS

DS and AJK conceived the study. DS planned and performed the wetlab experiments with guidance from AJK and experimental support from AV and ZD. DS and AJK wrote the paper.

## DECLARATION OF INTERESTS

There are no competing interests.

## STAR METHODS

### Cell culture and cerebral organoid generation

Feeder-independent H9 hESCs (WA09) were obtained from WiCell. Cells were maintained in 6-well tissue culture dishes (Fisher Scientific Corning Costar) coated with 0.5 mg/cm^2^ Vitronectin (VTN-N) (Thermo Fisher Scientific) in E8 medium (Thermo Fisher Scientific) and passaged using standard protocols. Angelman Syndrome (AS) iPSCs were developed by the laboratories of Stormy J. Chamberlain, PhD and Marc Lalande, PhD, University of Connecticut and obtained from Kerafast (Chamberlain et al., 2010). iPSCs were initially cultured on irradiated mouse embryonic fibroblasts (MEFs) with standard hESC medium: DMEM/F12 (Corning) supplied with 20 % KnockOut serum replacement (KOSR) (Thermo Fisher Scientific), 3 % fetal bovine serum (FBS) (VWR Life Science Seradigm), 1 % non-essential amino acids (NEAA) (HyClone), 1 % glutamax (Thermo Fisher Scientific), 4 μg/mL basic Fibroblast Growth Factor (Thermo Fisher Scientific 13256029) and 100 mM 2-mercaptoethanol (VWR), before being slowly adapted to E8 and vitronectin conditions. Cells were maintained in a humid incubator at 37 °C with 5 % (vol/vol) CO2. Cerebral organoids from H9 hESCs and AS iPSCs were generated and maintained using the same protocol as described (Lancaster and Knoblich, 2014; Lancaster et al., 2013).

### Topotecan and Indotecan treatment

Topotecan was purchased from Sigma Aldrich. Indotecan (LMP400) was obtained from the Developmental Therapeutics Program (DTP) branch, National Cancer Institute. Both drugs were reconstituted in dimethyl sulfoxide (DMSO) to 10 mM stock concentration, aliquoted to prevent freeze-thaw cycles, protected from light, and stored at − 20 °C. Drugs were directly added to AS cerebral organoids at 1 μM final concentration in culture medium. Organoids were cultured for 72 hrs without a fresh medium change to allow time for un-silencing of paternal *UBE3A*. For long term effects of single drug exposure experiments, the first drug-free medium change was performed 3-days after the single drug administration. Samples were collected 7 and 14 days after the fresh medium change (total of 10 and 17 days from initial drug exposure). For experiments testing the effects of drug exposure time, organoid culture medium was replaced daily with fresh medium containing drugs.

### Transfection of HEK 293FT Cells and Lentivirus Particles Production

To generate lentivirus particles, HEK 293FT cells (ThermoFisher) were seeded on a 6-well plate at a density of 5.0 × 10^6^ cells/ml in complete cultured media (DMEM with high glucose containing 10% FBS and non-essential amino acids; Corning). When cells reached 80% confluence, the medium was replaced with Opti-MEM reduced serum medium containing GlutaMax (ThermoFisher) and exposed to 25 μM chloroquine diphosphate (Sigma-Aldrich) for 5 hours before being transfected with the plasmid mixture. The plasmid mixture consisted of pLenti-CamKIIa-GFP (Addgene, Plasmid #96941), pCMVR8.74 (packaging plasmid, Addgene, Plasmid #22036), pCMV-VSV-G (envelope plasmid, Addgene, Plasmid #8454), and pAdVAntage vector (Promega, E1711) each at 300 fmol. Polyethyleneimine (PEI, Sigma) was used as the transfection reagent at a ratio of 3:1 (PEI:DNA). PEI was combined with the plasmid mixture, incubated for 20 minutes at room temperature, and spread drop-wise over the culture. A fresh media change was performed after 18 hours. Media containing lentiviral particles were harvested 48 and 72 hours post transfection, spun down at 500 g for 5 min and filtered using 0.45 μm PES syringe filters (VWR). The particles were concentrated by centrifugation at 2,500 g for 15 min using Amicon Ultra-15 centrifugal filter units (EMDMillipore, UFC910008). Concentrated lentivirus was aliquoted and stored at −80°C. hCOs were transduced by incubating 100μL pLenti-CamKIIa-GFP virus in 1mL cerebral organoid differentiation media for 12-hours.

### Histology and Immunofluorescence

Tissues were fixed in 4% paraformaldehyde for 15min at 4 °C followed by 3 × 10 minute PBS washes. Tissues were allowed to sink in 30 % sucrose overnight and then embedded in 10 % gelatin/7.5 % sucrose. Embedded tissues were frozen in an isopentane bath between −50 and −30 °C and stored at −80 °C. Frozen blocks were cryosectioned to 30-μm. For immunohistochemistry, sections were blocked and permeabilized in 1 % Triton X-100 and 5 % normal donkey serum in PBS. Sections were then incubated with primary antibodies in 0.3 % Triton X-100, 1 % normal donkey serum in PBS overnight at 4 °C in a humidity chamber. The primary antibody details and dilutions are listed in Table S1. Sections were then incubated with secondary antibodies in 0.3 % Triton X-100, 1 % normal donkey serum in PBS for 2h at RT, and nuclei were stained with DAPI (Invitrogen). Slides were mounted using ProLong Antifade Diamond (Thermo Fisher Scientific). Secondary antibodies used were donkey Alexa Fluor 488, 546 and 647 conjugates (Invitrogen, 1:500). Images were taken using a Nikon A1R confocal microscope (Nikon Instruments). High magnification images captured using thin (1.5 μm) optical sectioning. All samples within experiments were processed at the same time, imaged using the same microscope settings, and adjusted identically for quantification purposes. Quantifications were performed manually except for Figure 4C, where a CellProfiler pipeline automated the identification first of nucleic using DAPI, DAPI that were positive for GFP, and finally the mean background-subtracted UBE3A intensity or percent nuclei that had positive UBE3A signal above background. For quantification, intensities of all channels were maintained equally across all images. For displays, individual channels were balanced equally across the entire image.

### RNA extraction and qPCR

For each qPCR sample 6-12 hCOs were collected in 2mL RNAse-free tubes and chilled on ice throughout the procedure. hCOs were washed 3 times in cold PBS. Matrigel was dissolved by incubating the hCOs in chilled Cell Recovery Solution (Corning, cat. no. 354253) for 1h at 4 °C. The dissolved Matrigel was removed by rinsing 3 times in cold PBS. Total RNA was isolated using Direct-zol RNA MicroPrep Kit (Zymo Research) according to the manufacturer’s protocol. cDNA synthesis was performed using 900 ng of total RNA and the iScript Reverse Transcription Kit (BIO-RAD) according to the manufacturer’s protocol. qPCR reactions were performed using IQ Multiplex Powermix (BIO-RAD) on a BIO-RAD 384-well machine (CXF384) with PrimePCR probe assays (BIO-RAD). Unique assay id for *UBE3A* primers and probe is qHsaCIP0031486. Primer pairs and probes for *UBE3A-ATS* (RT-17 designed by Runte and colleagues) (Runte et al., 2001), *HPRT* and *TBP* were custom designed and are listed in Table S2. Individual primer pairs and probes were tested before multiplexing reactions. Analysis of *UBE3A* and *UBE3A-ATS* expression along with two reference genes *TBP* and *HPRT* was performed in triplicate using Excel by calculating the ΔΔCt value. Data are presented as expression level (2^−ΔΔCt^) relative to *TBP* or *HPRT*.

## References

Avagliano Trezza, R., Sonzogni, M., Bossuyt, S.N. V., Zampeta, F.I., Punt, A.M., van den Berg, M., Rotaru, D.C., Koene, L.M.C., Munshi, S.T., Stedehouder, J., et al. (2019). Loss of nuclear UBE3A causes electrophysiological and behavioral deficits in mice and is associated with Angelman syndrome. Nat. Neurosci. 1.

Bagley, J.A., Reumann, D., Bian, S., Lévi-Strauss, J., and Knoblich, J.A. (2017). Fused cerebral organoids model interactions between brain regions. Nat. Methods 14, 743–751.

Bailus, B.J., Pyles, B., McAlister, M.M., O’Geen, H., Lockwood, S.H., Adams, A.N., Nguyen, J.T.T., Yu, A., Berman, R.F., and Segal, D.J. (2016). Protein Delivery of an Artificial Transcription Factor Restores Widespread Ube3a Expression in an Angelman Syndrome Mouse Brain. Mol. Ther. 24, 548–555.

Bar, S., Schachter, M., Eldar-Geva, T., and Benvenisty, N. (2017). Large-Scale Analysis of Loss of Imprinting in Human Pluripotent Stem Cells. Cell Rep. 19, 957–968.

Bartolomei, M.S., and Ferguson-Smith, A.C. (2011). Mammalian genomic imprinting. Cold Spring Harb. Perspect. Biol. 3, 1–17.

Bershteyn, M., Nowakowski, T.J., Pollen, A.A., Di Lullo, E., Nene, A., Wynshaw-Boris, A., and Kriegstein, A.R. (2017). Human iPSC-Derived Cerebral Organoids Model Cellular Features of Lissencephaly and Reveal Prolonged Mitosis of Outer Radial Glia. Cell Stem Cell 20, 435–449.e4.

Birey, F., Andersen, J., Makinson, C.D., Islam, S., Wei, W., Huber, N., Fan, H.C., Metzler, K.R.C., Panagiotakos, G., Thom, N., et al. (2017). Assembly of functionally integrated human forebrain spheroids. Nature 545, 54–59.

Brockmann, K., Böhm, R., and Bürger, J. (2002). Exceptionally mild Angelman syndrome phenotype associated with an incomplete imprinting defect. J. Med. Genet. 39, e51.

Burette, A.C., Judson, M.C., Burette, S., Phend, K.D., Philpot, B.D., and Weinberg, R.J. (2017). Subcellular organization of UBE3A in neurons. J. Comp. Neurol. 525, 233–251.

Camp, J.G., Badsha, F., Florio, M., Kanton, S., Gerber, T., Wilsch-Bräuninger, M., Lewitus, E., Sykes, A., Hevers, W., Lancaster, M., et al. (2015). Human cerebral organoids recapitulate gene expression programs of fetal neocortex development. Proc. Natl. Acad. Sci. 201520760.

Chamberlain, S.J., and Brannan, C.I. (2001). The Prader–Willi Syndrome Imprinting Center Activates the Paternally Expressed Murine Ube3a Antisense Transcript but Represses Paternal Ube3a. Genomics 73, 316–322.

Chamberlain, S.J., Chen, P.-F., Ng, K.Y., Bourgois-Rocha, F., Lemtiri-Chlieh, F., Levine, E.S., and Lalande, M. (2010). Induced pluripotent stem cell models of the genomic imprinting disorders Angelman and Prader-Willi syndromes. Proc. Natl. Acad. Sci. U. S. A. 107, 17668–17673.

Choi, S.H., Kim, Y.H., Quinti, L., Tanzi, R.E., and Kim, D.Y. (2016). 3D culture models of Alzheimer’s disease: A road map to a “cure-in-a-dish.” Mol. Neurodegener. 11, 75.

Dang, J., Tiwari, S.K., Lichinchi, G., Qin, Y., Patil, V.S., Eroshkin, A.M., and Rana, T.M. (2016). Zika Virus Depletes Neural Progenitors in Human Cerebral Organoids through Activation of the Innate Immune Receptor TLR3. Cell Stem Cell 19, 258–265.

Dindot, S. V., Antalffy, B.A., Bhattacharjee, M.B., and Beaudet, A.L. (2008). The Angelman syndrome ubiquitin ligase localizes to the synapse and nucleus, and maternal deficiency results in abnormal dendritic spine morphology. Hum. Mol. Genet. 17, 111–118.

Englund, C. (2005). Pax6, Tbr2, and Tbr1 Are Expressed Sequentially by Radial Glia, Intermediate Progenitor Cells, and Postmitotic Neurons in Developing Neocortex. J. Neurosci. 25, 247–251.

Le Fevre, A., Beygo, J., Silveira, C., Kamien, B., Clayton-Smith, J., Colley, A., Buiting, K., and Dudding-Byth, T. (2017). Atypical Angelman syndrome due to a mosaic imprinting defect: Case reports and review of the literature. Am. J. Med. Genet. Part A 173, 753–757.

Fink, J.J., Robinson, T.M., Germain, N.D., Sirois, C.L., Bolduc, K.A., Ward, A.J., Rigo, F., Chamberlain, S.J., and Levine, E.S. (2017). Disrupted neuronal maturation in Angelman syndrome-derived induced pluripotent stem cells. Nat. Commun. 8, 15038.

Garcez, P.P., Loiola, E.C., Madeiro da Costa, R., Higa, L.M., Trindade, P., Delvecchio, R., Nascimento, J.M., Brindeiro, R., Tanuri, A., and Rehen, S.K. (2016). Zika virus impairs growth in human neurospheres and brain organoids. Science 352, 816–818.

Giandomenico, S.L., Mierau, S.B., Gibbons, G.M., Wenger, L.M.D., Masullo, L., Sit, T., Sutcliffe, M., Boulanger, J., Tripodi, M., Derivery, E., et al. (2019). Cerebral organoids at the air–liquid interface generate diverse nerve tracts with functional output. Nat. Neurosci. 22, 669–679.

Gonzalez-Gomez, M., and Meyer, G. (2014). Dynamic expression of calretinin in embryonic and early fetal human cortex. Front. Neuroanat. 8, 41.

Grier, M.D., Carson, R.P., and Lagrange, A.H. (2015). Toward a Broader View of Ube3a in a Mouse Model of Angelman Syndrome: Expression in Brain, Spinal Cord, Sciatic Nerve and Glial Cells. PLoS One 10, e0124649.

Gustin, R.M., Bichell, T.J., Bubser, M., Daily, J., Filonova, I., Mrelashvili, D., Deutch, A.Y., Colbran, R.J., Weeber, E.J., and Haas, K.F. (2010). Tissue-specific variation of Ube3a protein expression in rodents and in a mouse model of Angelman syndrome. Neurobiol. Dis. 39, 283–291.

Hillman, P.R., Christian, S.G.B., Doan, R., Cohen, N.D., Konganti, K., Douglas, K., Wang, X., Samollow, P.B., and Dindot, S. V (2017). Genomic imprinting does not reduce the dosage of UBE3A in neurons. Epigenetics and Chromatin 10, 27.

Hsiao, J.S., Germain, N.D., Wilderman, A., Stoddard, C., Wojenski, L.A., Villafano, G.J., Core, L., Cotney, J., and Chamberlain, S.J. (2019). A bipartite boundary element restricts UBE3A imprinting to mature neurons. Proc. Natl. Acad. Sci. 116, 2181–2186.

Huang, H.S., Allen, J.A., Mabb, A.M., King, I.F., Miriyala, J., Taylor-Blake, B., Sciaky, N., Dutton, J.W., Lee, H.M., Chen, X., et al. (2012). Topoisomerase inhibitors unsilence the dormant allele of Ube3a in neurons. Nature 481, 185–191.

Huang, X., Liu, J., Ketova, T., Fleming, J.T., Grover, V.K., Cooper, M.K., Litingtung, Y., and Chiang, C. (2010). Transventricular delivery of Sonic hedgehog is essential to cerebellar ventricular zone development. Proc. Natl. Acad. Sci. U. S. A. 107, 8422–8427.

Johansson, P.A., Irmler, M., Acampora, D., Beckers, J., and Götz, M. (2013). The transcription factor Otx2 regulates choroid plexus development and function. Development 140, 1055–1066.

Jones, K.A., Han, J.E., Debruyne, J.P., and Philpot, B.D. (2016). Persistent neuronal Ube3a expression in the suprachiasmatic nucleus of Angelman syndrome model mice. Sci. Rep. 6, 28238.

Judson, M.C., Sosa-Pagan, J.O., Del Cid, W.A., Han, J.E., and Philpot, B.D. (2014). Allelic specificity of Ube3a expression in the mouse brain during postnatal Development. J. Comp. Neurol. 522, 1874–1896.

Judson, M.C., Burette, A.C., Thaxton, C.L., Pribisko, A.L., Shen, M.D., Rumple, A.M., Del Cid, W.A., Paniagua, B., Styner, M., Weinberg, R.J., et al. (2017). Decreased Axon Caliber Underlies Loss of Fiber Tract Integrity, Disproportional Reductions in White Matter Volume, and Microcephaly in Angelman Syndrome Model Mice. J. Neurosci. 37, 7347–7361.

Kappil, M., Lambertini, L., and Chen, J. (2015). Environmental Influences on Genomic Imprinting. Curr. Environ. Heal. Reports 2, 155–162.

King, I.F., Yandava, C.N., Mabb, A.M., Hsiao, J.S., Huang, H.S., Pearson, B.L., Calabrese, J.M., Starmer, J., Parker, J.S., Magnuson, T., et al. (2013). Topoisomerases facilitate transcription of long genes linked to autism. Nature 501, 58–62.

Kishino, T., Lalande, M., and Wagstaff, J. (1997). UBE3A/E6-AP mutations cause Angelman syndrome. Nat. Genet. 15, 70–73.

Kummar, S., Chen, A., Gutierrez, M., Pfister, T.D., Wang, L., Redon, C., Bonner, W.M., Yutzy, W., Zhang, Y., Kinders, R.J., et al. (2016). Clinical and pharmacologic evaluation of two dosing schedules of indotecan (LMP400), a novel indenoisoquinoline, in patients with advanced solid tumors. Cancer Chemother. Pharmacol. 78, 73–81.

Lancaster, M.A., and Knoblich, J.A. (2014). Generation of cerebral organoids from human pluripotent stem cells. Nat. Protoc. 9, 2329–2340.

Lancaster, M.A., Renner, M., Martin, C.-A., Wenzel, D., Bicknell, L.S., Hurles, M.E., Homfray, T., Penninger, J.M., Jackson, A.P., and Knoblich, J.A. (2013). Cerebral organoids model human brain development and microcephaly. Nature 501.

LaSalle, J.M., Reiter, L.T., and Chamberlain, S.J. (2015). Epigenetic regulation of UBE3A and roles in human neurodevelopmental disorders. Epigenomics 7, 1213–1228.

Lee, H.M., Clark, E.P., Kuijer, M.B., Cushman, M., Pommier, Y., and Philpot, B.D. (2018). Characterization and structure-activity relationships of indenoisoquinoline-derived topoisomerase i inhibitors in unsilencing the dormant Ube3a gene associated with Angelman syndrome. Mol. Autism 9, 45.

Li, Y., Muffat, J., Omer, A., Bosch, I., Lancaster, M.A., Sur, M., Gehrke, L., Knoblich, J.A., and Jaenisch, R. (2017). Induction of Expansion and Folding in Human Cerebral Organoids. Cell Stem Cell 20, 385–396.e3.

Litzky, J.F., Deyssenroth, M.A., Everson, T.M., Lester, B.M., Lambertini, L., Chen, J., and Marsit, C.J. (2018). Prenatal exposure to maternal depression and anxiety on imprinted gene expression in placenta and infant neurodevelopment and growth. Pediatr. Res. 83, 1075–1083.

Lopez, S.J., Segal, D.J., and LaSalle, J.M. (2019). UBE3A: An E3 Ubiquitin Ligase With Genome-Wide Impact in Neurodevelopmental Disease. Front. Mol. Neurosci. 11, 476.

Di Lullo, E., and Kriegstein, A.R. (2017). The use of brain organoids to investigate neural development and disease. Nat. Rev. Neurosci. 18, 573–584.

Luo, C., Lancaster, M.A., Castanon, R., Nery, J.R., Knoblich, J.A., and Ecker, J.R. (2016). Cerebral Organoids Recapitulate Epigenomic Signatures of the Human Fetal Brain. Cell Rep. 17, 3369–3384.

Mabb, A.M., Judson, M.C., Zylka, M.J., and Philpot, B.D. (2011). Angelman syndrome: Insights into genomic imprinting and neurodevelopmental phenotypes. Trends Neurosci. 34, 293–303.

Mariani, J., Coppola, G., Zhang, P., Abyzov, A., Provini, L., Tomasini, L., Amenduni, M., Szekely, A., Palejev, D., Wilson, M., et al. (2015). FOXG1-Dependent Dysregulation of GABA/Glutamate Neuron Differentiation in Autism Spectrum Disorders. Cell 162, 375–390.

Mason, J.O., and Price, D.J. (2016). Building brains in a dish: Prospects for growing cerebral organoids from stem cells. Neuroscience 334, 105–118.

Meng, L., Person, R.E., and Beaudet, A.L. (2012). Ube3a-ATS is an atypical RNA polymerase II transcript that represses the paternal expression of Ube3a. Hum. Mol. Genet. 21, 3001–3012.

Meng, L., Person, R.E., Huang, W., Zhu, P.J., Costa-Mattioli, M., and Beaudet, A.L. (2013). Truncation of Ube3a-ATS Unsilences Paternal Ube3a and Ameliorates Behavioral Defects in the Angelman Syndrome Mouse Model. PLoS Genet. 9, e1004039.

Meng, L., Ward, A.J., Chun, S., Bennett, C.F., Beaudet, A.L., and Rigo, F. (2014). Towards a therapy for Angelman syndrome by targeting a long non-coding RNA. Nature 518, 409–412.

Miao, S., Chen, R., Ye, J., Tan, G.-H., Li, S., Zhang, J., Jiang, Y.-h., and Xiong, Z.-Q. (2013). The Angelman Syndrome Protein Ube3a Is Required for Polarized Dendrite Morphogenesis in Pyramidal Neurons. J. Neurosci. 33, 327–333.

Pasca, A.M., Sloan, S.A., Clarke, L.E., Tian, Y., Makinson, C.D., Huber, N., Kim, C.H., Park, J.Y., O’Rourke, N.A., Nguyen, K.D., et al. (2015). Functional cortical neurons and astrocytes from human pluripotent stem cells in 3D culture. Nat. Methods 12, 671–678.

Qian, X., Nguyen, H.N., Song, M.M., Hadiono, C., Ogden, S.C., Hammack, C., Yao, B., Hamersky, G.R., Jacob, F., Zhong, C., et al. (2016). Brain-Region-Specific Organoids Using Mini-bioreactors for Modeling ZIKV Exposure. Cell 165, 1238–1254.

Quadrato, G., Nguyen, T., Macosko, E.Z., Sherwood, J.L., Min Yang, S., Berger, D.R., Maria, N., Scholvin, J., Goldman, M., Kinney, J.P., et al. (2017). Cell diversity and network dynamics in photosensitive human brain organoids. Nature 545, 48–53.

Raja, W.K., Mungenast, A.E., Lin, Y.-T., Ko, T., Abdurrob, F., Seo, J., and Tsai, L.-H. (2016). Self-Organizing 3D Human Neural Tissue Derived from Induced Pluripotent Stem Cells Recapitulate Alzheimer’s Disease Phenotypes. PLoS One 11, e0161969.

Rotaru, D.C., van Woerden, G.M., Wallaard, I., and Elgersma, Y. (2018). Adult Ube3a Gene Reinstatement Restores the Electrophysiological Deficits of Prefrontal Cortex Layer 5 Neurons in a Mouse Model of Angelman Syndrome. J. Neurosci. 38, 8011–8030.

Rougeulle, C., Glatt, H., and Lalande, M. (1997). The Angelman syndrome candidate gene, UBE3A/E6-AP, is imprinted in brain. Nat. Genet. 17, 14–15.

Runte, M., Hüttenhofer, A., Groß, S., Kiefmann, M., Horsthemke, B., and Buiting, K. (2001). The IC-SNURF–SNRPN transcript serves as a host for multiple small nucleolar RNA species and as an antisense RNA for UBE3A. Hum. Mol. Genet. 10, 2687–2700.

Sadhwani, A., Sanjana, N.E., Willen, J.M., Calculator, S.N., Black, E.D., Bean, L.J.H., Li, H., and Tan, W.-H. (2018). Two Angelman families with unusually advanced neurodevelopment carry a start codon variant in the most highly expressed *UBE3A* isoform. Am. J. Med. Genet. Part A 176, 1641–1647.

Saito, T., Hanai, S., Takashima, S., Nakagawa, E., Okazaki, S., Inoue, T., Miyata, R., Hoshino, K., Akashi, T., Sasaki, M., et al. (2011). Neocortical layer formation of human developing brains and lissencephalies: Consideration of layer-specific marker expression. Cereb. Cortex 21, 588–596.

Sato, M., and Stryker, M.P. (2010). Genomic imprinting of experience-dependent cortical plasticity by the ubiquitin ligase gene Ube3a. Proc. Natl. Acad. Sci. 107, 5611–5616.

Silva-Santos, S., Woerden, G.M. van, Bruinsma, C.F., Mientjes, E., Jolfaei, M.A., Distel, B., Kushner, S.A., and Elgersma, Y. (2015). Ube3a reinstatement identifies distinct developmental windows in a murine Angelman syndrome model. J. Clin. Invest. 125, 2069–2076.

Singhmar, P., and Kumar, A. (2011). Angelman Syndrome Protein UBE3A Interacts with Primary Microcephaly Protein ASPM, Localizes to Centrosomes and Regulates Chromosome Segregation. PLoS One 6, e20397.

Sonzogni, M., Hakonen, J., Bernabé Kleijn, M., Silva-Santos, S., Judson, M.C., Philpot, B.D., van Woerden, G.M., and Elgersma, Y. (2019). Delayed loss of UBE3A reduces the expression of Angelman syndrome-associated phenotypes. Mol. Autism 10, 23.

Stanurova, J., Neureiter, A., Hiber, M., De Oliveira Kessler, H., Stolp, K., Goetzke, R., Klein, D., Bankfalvi, A., Klump, H., and Steenpass, L. (2016). Angelman syndrome-derived neurons display late onset of paternal UBE3A silencing. Sci. Rep. 6.

Vatsa, N., and Jana, N.R. (2018). UBE3A and Its Link With Autism. Front. Mol. Neurosci. 11, 448.

Velasco, S., Kedaigle, A.J., Simmons, S.K., Nash, A., Rocha, M., Quadrato, G., Paulsen, B., Nguyen, L., Adiconis, X., Regev, A., et al. (2019). Individual brain organoids reproducibly form cell diversity of the human cerebral cortex. Nature 570, 523–527.

Xiang, Y., Tanaka, Y., Patterson, B., Kang, Y.-J., Govindaiah, G., Roselaar, N., Cakir, B., Kim, K.-Y., Lombroso, A.P., Hwang, S.-M., et al. (2017). Fusion of Regionally Specified hPSC-Derived Organoids Models Human Brain Development and Interneuron Migration. Cell Stem Cell 21, 383–398.e7.

Yamasaki, K., Joh, K., Ohta, T., Masuzaki, H., Ishimaru, T., Mukai, T., Niikawa, N., Ogawa, M., Wagstaff, J., and Kishino, T. (2003). Neurons but not glial cells show reciprocal imprinting of sense and antisense transcripts of Ube3a. Hum. Mol. Genet. 12, 837–847.

Yi, J.J., Paranjape, S.R., Walker, M.P., Choudhury, R., Wolter, J.M., Fragola, G., Emanuele, M.J., Major, M.B., and Zylka, M.J. (2017). The autism-linked UBE3A T485A mutant E3 ubiquitin ligase activates the Wnt/β-catenin pathway by inhibiting the proteasome. J. Biol. Chem. 292, 12503–12515.

