## Supplemental Information for "Human cerebral organoids capture the spatiotemporal complexity and disease dynamics of UBE3A"

### SUPPLEMENTARY FIGURES

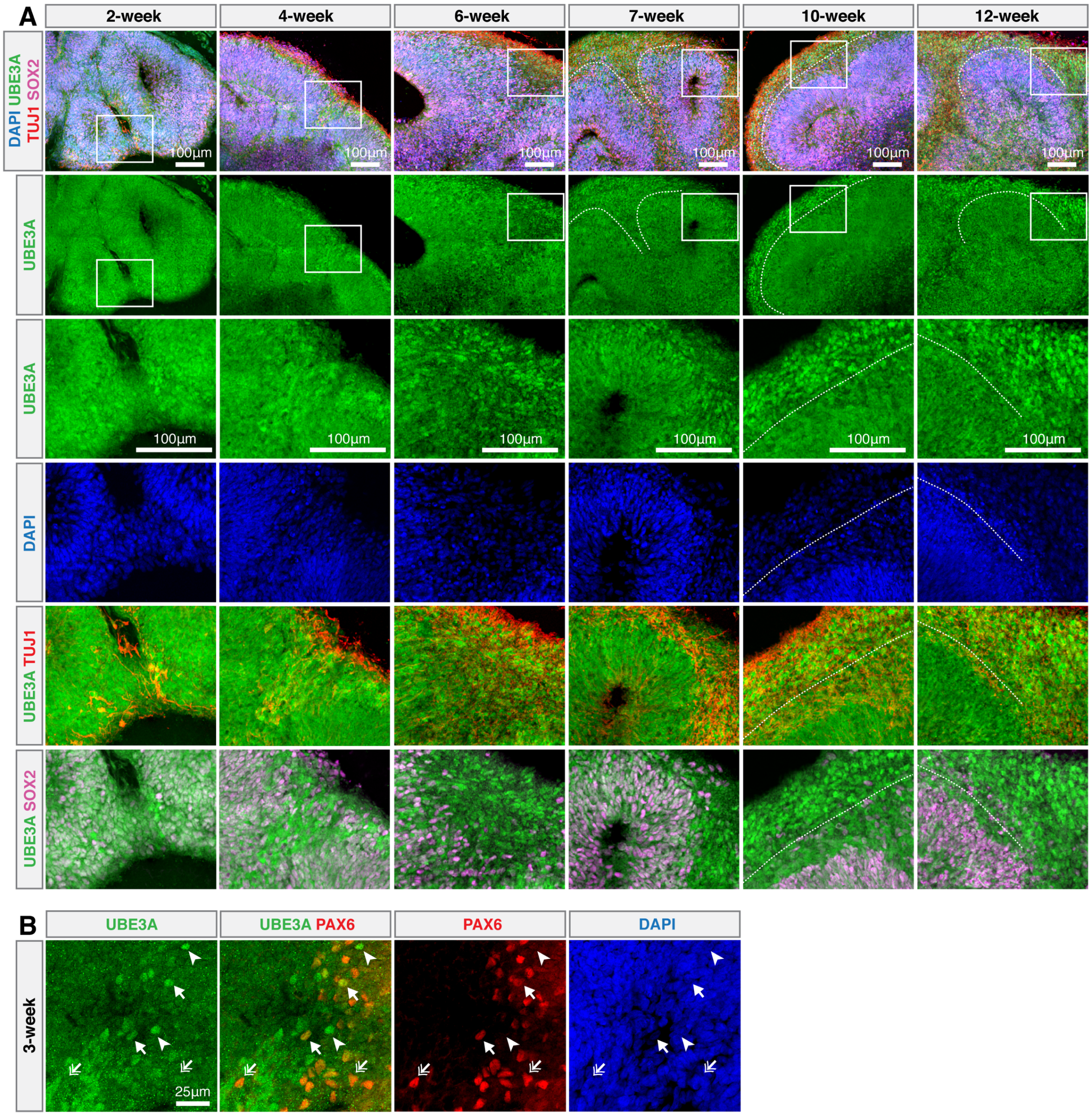


**Figure S1 related to Figure 1. Additional time-points in hCO development revealing the cytoplasmic to nuclear transition of UBE3A**

(A) Immunostaining time course of neurotypical hCO neurodevelopment. (A) Boxes bound high magnification images. Nuclear UBE3A in neurons (arrows). Diffuse UBE3A in SOX2+ cells (arrow heads). Decreasing cytoplasmic UBE3A over time (double arrows). Dotted white lines delineate boundaries between TUJ1+ and SOX2+ cells.

(B) Nuclear (arrows) and diffuse (double arrows) UBE3A in PAX6+ cells. Nuclear UBE3A in PAX6-/weak cells (arrow heads).

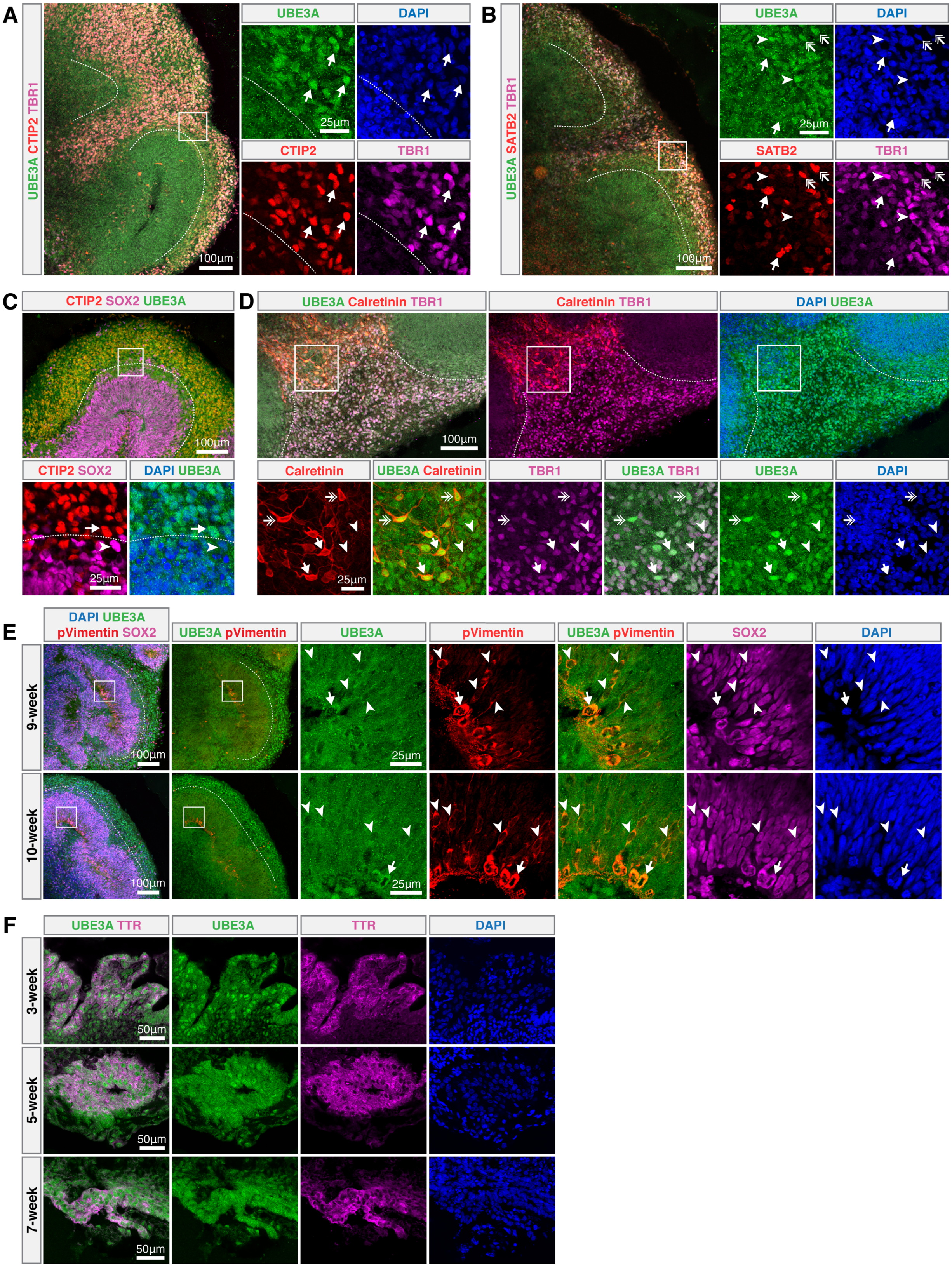
A
UBE3A CTIP2 TBR1
UBE3A
DAPI
CTIP2
TBR1
25µm
100µm
B
UBE3A SATB2 TBR1
UBE3A
DAPI
SATB2
TBR1
25µm
100µm
C
CTIP2 SOX2 UBE3A
100µm
CTIP2 SOX2
DAPI UBE3A
25µm
D
UBE3A Calretinin TBR1
Calretinin TBR1
DAPI UBE3A
100µm
Calretinin
UBE3A Calretinin
TBR1
UBE3A TBR1
UBE3A
DAPI
25µm
E
DAPI UBE3A
pVimentin SOX2
UBE3A pVimentin
UBE3A
pVimentin
UBE3A pVimentin
SOX2
DAPI
9-week
10-week
100µm
25µm
F
UBE3A TTR
UBE3A
TTR
DAPI
3-week
5-week
7-week
50µm

**Figure S2 related to Figure 2. UBE3A in cerebral cortex-like, progenitor, and choroid plexus-like compartments.**

UBE3A in (A-D) cortical cells, (E) radial glia, and (F) choroid plexus in neurotypical hCOs. Boxes bound high magnification images.

(A, arrows) CTIP2, TBR1 and nuclear UBE3A co-localize.

(B) Nuclear UBE3A in (double arrows) TBR1+/SATB2+ and (arrow heads) TBR+/SATB2- cells. Diffuse UBE3A in (arrows) TBR1-/SATB2+ cells.

(C) Nuclear UBE3A in CTIP2+ neurons (arrow). Diffuse UBE3A in SOX2+ progenitors (arrow head).

(D) Nuclear UBE3A in TBR1+/Calretinin+ (arrows), TBR1weak/Calretinin+ (double arrows) and TBR+/Calretinin- (arrow heads) cells.

(E) Relative absence of UBE3A from nuclei of pVimentin+/SOX2+ cells (arrow heads). Higher intensity UBE3A excluded from condensed chromatin in actively dividing radial glia (arrows).

(F) Nuclear UBE3A weakens with time in TTR+ cells.

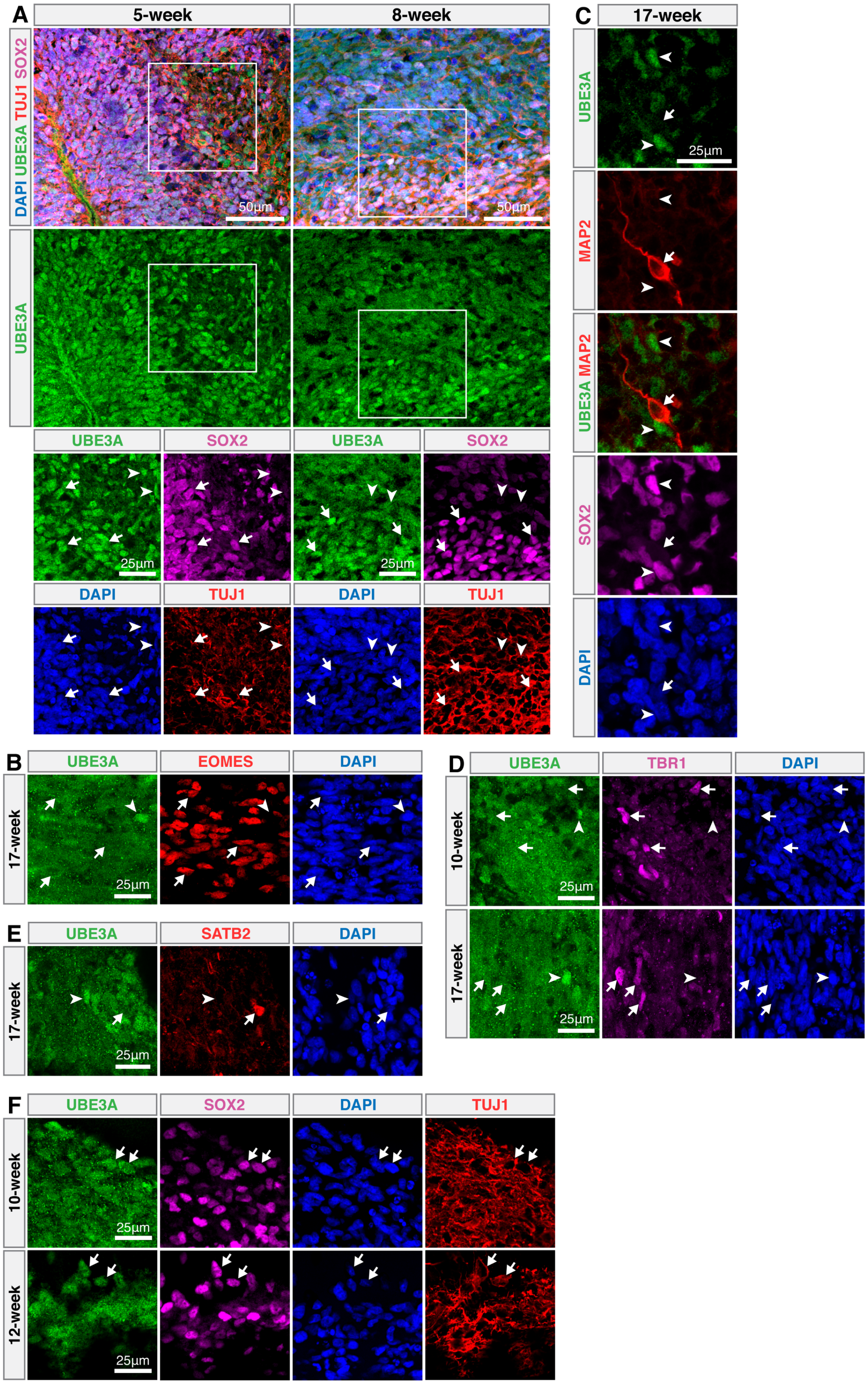
A
5-week
8-week
DAPI UBE3A TUJ1 SOX2
UBE3A
50µm
50µm
UBE3A
SOX2
UBE3A
SOX2
25µm
25µm
DAPI
TUJ1
DAPI
TUJ1
C
17-week
UBE3A
25µm
MAP2
UBE3A MAP2
SOX2
DAPI
B
UBE3A
EOMES
DAPI
17-week
25µm
D
UBE3A
TBR1
DAPI
10-week
17-week
25µm
25µm
E
UBE3A
SATB2
DAPI
17-week
25µm
F
UBE3A
SOX2
DAPI
TUJ1
10-week
12-week
25µm
25µm

**Figure S3 related to Figure 3. Paternal UBE3A expression and localization during AS hCO development.**

(A) Nuclear UBE3A in 5-8 week old SOX2+ progenitors (arrows). Nuclear UBE3A in TUJ1+/SOX2- neurons at 5 weeks becomes diffuse at 8 weeks (arrow heads).

(B) Weak/diffuse UBE3A in EOMES+ cells (arrows). Nuclear UBE3A in some EOMES- cells (arrow head).

(C) UBE3A absent in MAP2+/SOX2- neurons (arrow). Paternal UBE3A expressed in SOX2+ progenitors after 17 weeks (arrow heads).

(D) Weak/diffuse UBE3A in TBR1+ cells (arrows). Nuclear UBE3A in some TBR1- cells (arrow heads).

(E) Weak/diffuse UBE3A in SATB2+ cells (arrow). Nuclear UBE3A in some SATB2- cells (arrow head).

(F) Nuclear UBE3A in SOX2+/TUJ1+ immature neurons (arrows).

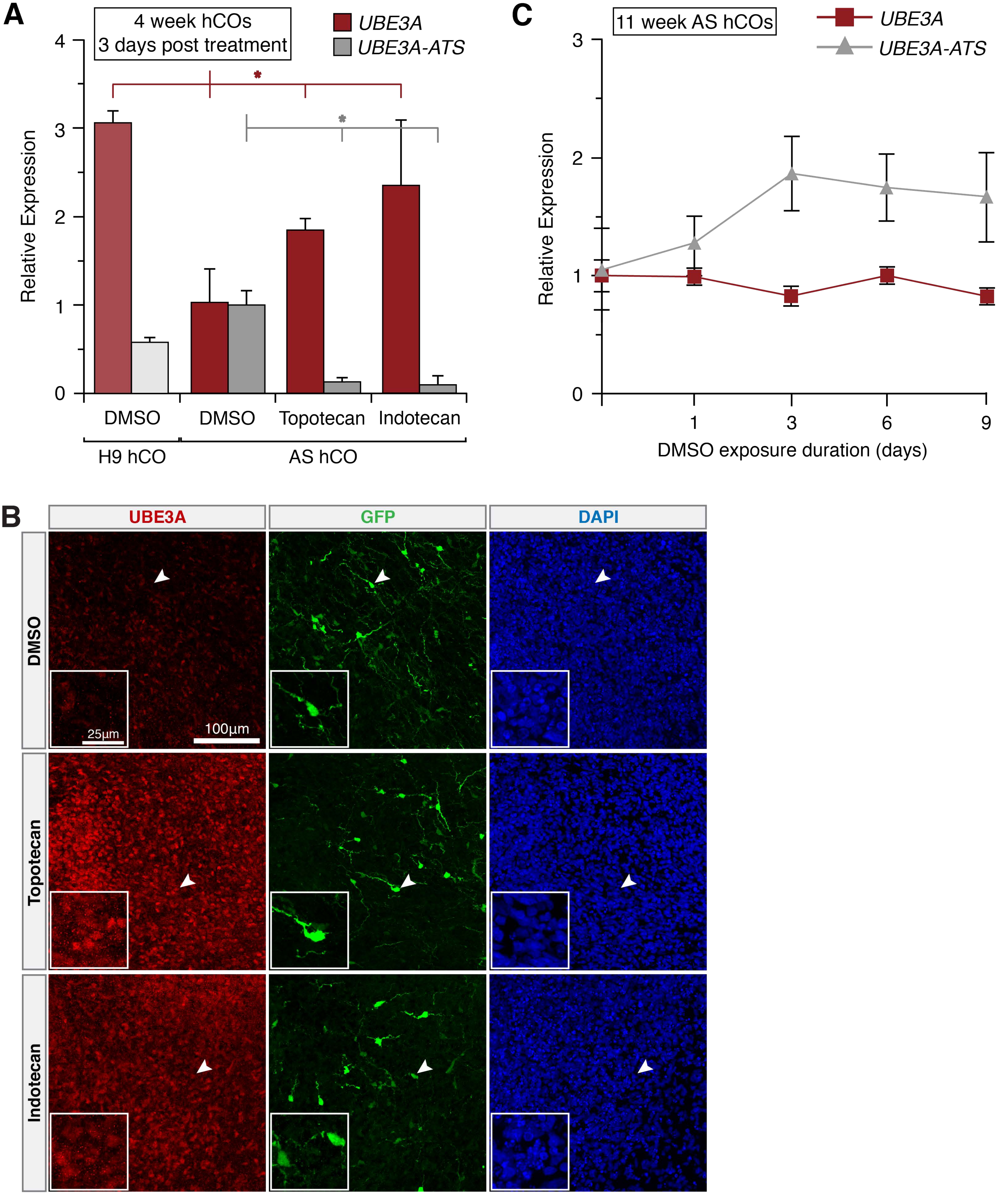

A
4 week hCOs
3 days post treatment
UBE3A
UBE3A-ATS
Relative Expression
4
3
2
1
0
DMSO
DMSO
Topotecan
Indotecan
H9 hCO
AS hCO
C
11 week AS hCOs
UBE3A
UBE3A-ATS
Relative Expression
3
2
1
0
1
3
6
9
DMSO exposure duration (days)
B
UBE3A
GFP
DAPI
DMSO
Topotecan
Indotecan
25µm
100µm

**Figure S4 related to Figure 4. Paternal *UBE3A* can be pharmacologically re-activated in AS hCOs.**

(A) *UBE3A* and *UBE3A-ATS* expression in 15 week old AS hCOs 3 days after a single drug treatment.

(B) Additional immunostained images of 11 week old AS hCOs with CamKIIa-GFP neurons. Insets zoom in on arrow heads.

(C) *UBE3A* and *UBE3A-ATS* expression after 1-9-days of continuous vehicle (DMSO) treatment in 11 week old AS hCOs.

### SUPPLEMENTARY TABLES

#### Table S1. Primary antibody information

| Antigen | Host | Supplier | Cat. No. | RRID | Dilution |
|---|---|---|---|---|---|
| SOX2 | Goat | R&D systems | AF2018 | AB_355110 | 1:20 |
| TUJ1 | Mouse | Sigma Aldrich | T8578 | AB_1841228 | 1:100 |
| UBE3A | Rabbit | Bethyl Laboratories | A300-351A | AB_185563 | 1:250 |
| pVimentin | Mouse | MBL international | D076-3S | AB_592962 | 1:200 |
| MAP2 | Mouse | Sigma Aldrich | M1406 | AB_477171 | 1:250 |
| TTR | Sheep | BIO-RAD | AHP1837 | AB_2212089 | 1:100 |
| TBR1 | Chicken | Sigma Aldrich | WH0010716M1 | AB_1843877 | 1:100 |
| EOMES | Mouse | R&D systems | MAB6166 | AB_10919889 | 1:25 |
| CTIP2 | Rat | abcam | ab18465 | AB_2064130 | 1:100 |
| SATB2 | Mouse | abcam | ab51502 | AB_882455 | 1:100 |
| Calretinin | Mouse | Millipore Sigma | MAB1568 | AB_94259 | 1:100 |
| GFP | Chicken | Abcam | ab13970 | AB_300798 | 1:500 |

#### Table S2. RT-qPCR primer information

| Target | Gene ID | Forward primer | Reverse primer | Probe | Amplicon size |
|---|---|---|---|---|---|
| *UBE3A-ATS* | 104472715 | GGCACTGAAAAT GTGGCATCCAG | GGTGTGTCAGCT GTGCTGGTGTC | AGCCAAAGAGTACTC TTCCTCAGTCATCCT | 120 |
| *TBP* | 6908 | GGGCACCACTCC ACTGTATC | CGAAGTGCAATG GTCTTTAGG | ATGACTCCCATGACCC CCATCACTCCT | 100 |
| *HPRT* | 3251 | TGACACTGGCAA AACAATGCA | GGTCCTTTTCACC AGCAAGCT | TGCTTTCCTTGGTCAG GCAGTATAATCCA | 94 |
